# Beamforming seizures from the temporal lobe

**DOI:** 10.1101/2021.11.19.469291

**Authors:** Luis Garcia Dominguez, Apameh Tarazi, Taufik Valiante, Richard Wennberg

## Abstract

**Background:** Surgical treatment of drug-resistant temporal lobe epilepsy (TLE) depends on proper identification of the seizure onset zone (SOZ), and differentiation of mesial, temporolimbic seizure onsets from temporal neocortical seizure onsets. Non-invasive source imaging using electroencephalography (EEG) and magnetoencephalography (MEG) can provide accurate information on interictal spike localization; however, EEG and MEG have low sensitivity for epileptiform activity restricted to deep temporolimbic structures. Moreover, in mesial temporal lobe epilepsy (MTLE), interictal spikes frequently arise in neocortical foci distant from the SOZ, rendering interictal spike localization potentially misleading for presurgical planning.

**Methods:** In this study, we used two different beamformer techniques applied to the MEG signal of ictal events acquired during EEG-MEG recordings in six patients with TLE (three neocortical, three MTLE). The ictal source localization results were compared to the patients’ ground truth SOZ localizations determined from intracranial EEG and/or clinical, neuroimaging and postsurgical outcome evidence.

**Results:** Beamformer analysis proved to be highly accurate in all cases and able to reliably identify focal seizure onsets localized to mesial, temporolimbic structures. In three patients, interictal spikes were either absent, too complex for inverse dipole modeling, or localized to anterolateral temporal neocortex distant to a mesial temporal SOZ.

**Conclusions:** This report demonstrates the suitability of MEG beamformer analysis of ictal events in TLE, which can supersede or complement the traditional analysis of interictal spikes. The method outlined is applicable to any type of epileptiform event, greatly expanding the information value of MEG and broadening its utility for presurgical recording in epilepsy.

## Introduction

Surgical treatment of drug-resistant temporal lobe epilepsy (TLE) depends on proper identification of the seizure onset zone (SOZ), and on the differentiation of mesial, temporolimbic seizure onsets from temporal neocortical seizure onsets. This presents a challenge for non-invasive presurgical investigations using scalp electroencephalography (EEG) or magnetoencephalography (MEG), especially in mesial temporal lobe epilepsy (MTLE), where the anatomically and physiologically unique temporolimbic structures prone to seizure generation are situated deep in the brain, where epileptiform activity is often “invisible” to EEG and MEG.

The definitive gold standard for identification of the SOZ is intracranial EEG (iEEG), provided that the implanted electrodes are located in close proximity to the SOZ. In practice, intracranial and extracranial recording modalities can work together, with non-invasive source localization helping to guide iEEG electrode placement. In some cases, when source localization is concordant with clinical and neuroimaging information suggesting a discrete, focal seizure generator, iEEG can be bypassed and a patient may proceed directly to surgery. In such cases, a postsurgical seizure-free outcome is taken as evidence that the “epileptogenic zone” was correctly identified. The epileptogenic zone, by definition, includes the SOZ, but it is not necessarily limited to the SOZ, making it a less good standard against which to assess the accuracy of EEG or MEG ictal source localization. Nevertheless, a postsurgical seizure-free outcome, combined with a strong presurgical hypothesis on location of the SOZ, can serve as a reasonable surrogate for ground truth in the absence of iEEG.

Among many source localization methodologies, the equivalent current dipole (ECD) inverse modeling method, applied to interictal spikes, has been used most extensively for presurgical planning in TLE.^1–3^ However, interictal spikes may be dissociated from the SOZ, a situation especially common in MTLE.^2–4^ Moreover, it is often the case in MTLE that more than one area in the temporal lobe produces independent interictal spikes,^4^ limiting the utility of interictal spike source localization for determining whether a TLE patient’s seizures are of mesial or neocortical origin.^5^ For the localization of ictal events, the ECD model is limited in its application to seizures with sharp, discernible discharges, whereas most ictal events, as well as shorter “ictal-like” or polyspike events, show a multi-frequency, oscillatory pattern. In a recent systematic review and meta-analysis of interictal and ictal EEG and MEG source imaging studies it was found that various techniques demonstrated reasonably high accuracy in localizing epileptic foci, at least to a lobar or sublobar level, however, ictal MEG analyses, as performed to date, were noted to have the lowest specificity and accuracy.^6^

Notwithstanding, one would prefer to localize directly the seizure instead of the interictal spike, especially in MTLE.^4,5^ Furthermore, for the purposes of presurgical planning in patients with drug-resistant epilepsy, discrete localization to a particular gyrus/sulcus is the desired ideal, rather than simple identification of the correct lobe or broad sublobar region.^2^ What is needed is a sufficiently robust method that can be applied to any pattern of temporal dynamics, and that can also uncover ictal events occurring in deep structures. MEG is, in principle, a good candidate for capturing focal activity due to its low sensitivity to secondary currents, however, the ictal MEG literature is sparse.

Most ictal MEG studies have used the ECD model, alone or in addition to frequency-based methods,^7–14^ including a large study of 23 patients with extratemporal epilepsy who underwent both iEEG and surgical resection of the presumptive epileptogenic zone, which reported 70-75% accuracy for ictal localization at the level of lobar surface planes.^14^

A few studies have followed a different path, using distributed source models^15–18^ or beamforming^19^ to analyze ictal MEG recordings.^20^ A very large study of 44 patients with neocortical epilepsy concluded that extended-source minimum-norm estimation on seizure-specific frequency bands was more suitable for localization of ictal rhythms than single-point (ECD) solutions, but that modeling of deeply seated foci presented a challenge.^17^ A study of 13 patients with neocortical epilepsy reported 90% sublobar concordance between the SOZ localized by wavelet-based maximum entropy on the mean analysis of ictal MEG and that identified by iEEG or suggested by MRI.^18^ Badier et al.^19^ used a beamforming technique, linearly constrained minimum variance (LCMV), to analyze rhythmic ictal MEG patterns in six patients (five extratemporal, one MTLE) and found LCMV superior to the ECD model in terms of concordance with iEEG findings, but unable to resolve the deep hippocampal SOZ in their MTLE patient.

The use of beamformers for the spatial analysis of cortical rhythms is pervasive in cognitive neuroscience, however, in the assessment of epileptic foci beamforming has been mainly restricted to the recovery of virtual channels, typically obtained via LCMV^21^ or synthetic aperture magnetometry,^22^ aiming at some second step of feature evaluation. Most popular is the so-called kurtosis beamformer,^22–25^ in which kurtosis is evaluated on the virtual channels across interictal periods. Considering the rhythmicity that characterizes ictal events it is perhaps surprising that little work has been done to investigate the use of a frequency domain beamformer, such as dynamic imaging of coherent sources (DICS),^26–28^ as a potential tool for non-invasive presurgical localization of the SOZ.

In this paper, we present a beamforming method applied to the MEG signal of ictal events recorded in a series of six TLE patients, three with neocortical epilepsy and three with MTLE. We have chosen to focus in detail on this small group of patients, drawn from a large pool of patients with temporal and extratemporal epilepsy in whom we have recorded ictal events using MEG, because of the unique neuroanatomical and neurophysiological characteristics of MTLE, which merit special consideration in studies of epilepsy surgery,^4,5^ and because ground truth evidence of SOZ location is available for this group, against which the accuracy of the beamforming method can be directly compared.

## Methods

### Recording

MEG was acquired using an Elekta Neuromag TRIUX 306-channel system (Helsinki, Finland) with simultaneous 32-channel EEG. Patients were sleep-deprived prior to the 90-minute recording. Sampling frequency was 1000Hz; online filter bandwidth 0.1–330Hz. Artifact suppression used the default parameters of the spatiotemporal signal space separation algorithm implemented within the Elekta Maxfilter system (10-second time window, subspace correlation 0.980).

### Anatomy

Head position inside the MEG sensor array was determined using a head position indicator (HPI) with five coils attached to the scalp. A Polhemus 3D-Fastrak system (Colchester, USA) was used to digitize the head shape and HPI coil positions.

Individual T1-weighted anatomical MRI scans and the corresponding digitized head shapes were co-registered, initially manually, using anatomical landmarks, and then automatically, using an iterative closest point algorithm, as described previously.^29,30^ The lead field was computed using a realistically-shaped single-shell approximation.^31^ The source space comprised a regular 3D grid, with 5mm distance between adjacent nodes, inside the inner surface of the skull.

### Signal processing of ictal events

Ictal events were manually identified by a clinical neurophysiologist (RW) in the MEG-EEG recording, and used to epoch the ictal MEG signal into trials, typically 1-second long, with 50% overlap. Baseline (interictal) periods were visually selected and epoched with the same duration. Baseline segments were chosen to ensure the subject was in the same state of sleep or wakefulness as during the ictal events to be modeled. An arbitrarily large number (n=500) of these interictal epochs were randomly selected. Then, both ictal and interictal epochs were inspected for artifacts using different statistics such as variance, range, and kurtosis, and trials removed if they were visually considered outliers for any of these statistics. The power spectra of the two sets of epochs were then compared as the ratio, in decibels, of the average power over all channels. This ratio, for individual channels, is then visualized over schematic head diagrams (we will call them “topoplots”). In these plots, each pair of orthogonal planar gradiometers corresponding to the same sensor are averaged together. This is essential to enable detection of focal patterns in sensor space, which cannot be achieved by mapping the field from magnetometers.

Both elements, the relative power spectrum and the topoplots, help to determine the frequency of interest. Here the goal is to identify not only the frequency with the largest signal-to-noise ratio, but also one that displays a pattern that indicates focality, that is, high values restricted to a small neighbourhood in sensor space. In some cases, the same focal pattern appears across all or a wide range of frequencies; in other cases, the pattern can morph over the frequency domain, displaying two or more distinct configurations. In the latter scenario, all of the main frequency peaks for the distinct patterns are analyzed independently.

We follow the principle of common filters where both sets of trials are pooled together in order to obtain a common beamformer filter. This filter is then applied independently to each set of trials. The end result is displayed as the contrast between both conditions, expressed as the relative change of power of ictal versus interictal periods over every node. In this report we will refer to this unitless entity as “beamformer power contrast” (BPC). The actual implementation of the DICS and LCMV beamformers, and the frequency analysis was done with the help of Fieldtrip.^32^ When evaluating the beamformer solution there is sometimes more than one local BPC maximum. If this is the case, and if the additional maxima are relevant in terms of the magnitude of the BPC, then those locations are also reported.

This method places an emphasis on beamforming each prevalent ictal frequency using DICS and/or bandpassed LCMV. If no single dominant frequency is present, unfiltered LCMV is also used. It is to be stressed that, although the cases are processed within a single conceptual pipeline, the parameters of the computation are specific to how an individual seizure manifests in its frequency and spatial domains.

### Source localization of interictal spikes

Interictal spikes were manually identified (RW) in the MEG-EEG recording and electromagnetic source imaging (EMSI) of interictal spike foci performed using CURRY 6 (Compumedics, Abbotsford, Australia), as described previously.^2,3,29^

### Patients

Cases were selected from 300 consecutive MEG-EEG recordings performed between 2015 and 2021 at the Mitchell Goldhar MEG Unit, Toronto Western Hospital, as part of the presurgical investigation of patients with drug-resistant epilepsy, and included all patients with temporal lobe ictal events who also had iEEG or other ground truth SOZ localization. Among the 300 patients, 75 patients had ictal events recorded with MEG, either clinical or subclinical seizures or shorter, ictal-like or polyspike bursts: 57 with extratemporal epilepsy and 18 with TLE. Ictal MEG beamformer localization in the extratemporal epilepsy group will be the subject of a different study. For this study, 12 of the TLE patients were excluded due to lack of definitive SOZ identification (i.e., neither iEEG nor clinical, MRI and postsurgical outcome evidence of unilateral MTLE), leaving six patients who met inclusion criteria. The characteristics of the six included patients are presented in Table 1. The three patients with neocortical TLE all had iEEG recording. Of the three MTLE patients, only the patient with normal brain MRI underwent iEEG; for the other two patients, both with unilateral hippocampal sclerosis, a postsurgical seizure-free outcome served, along with the clinical and MRI information, as the ground truth.

**Table 1.** Patient characteristics and ictal beamformer localizations.

| Patient | Age (years) | Epilepsy duration (years) | Brain MRI | iEEG | Distance from main ictal iEEG contacts to ictal beamformer solution (cm) |  | Distance between interictal ECD and ictal beamformer solutions (cm) | Surgical outcome 1-year (Engel) |
| --- | --- | --- | --- | --- | --- | --- | --- | --- |
| Mesial temporal lobe epilepsy |  |  |  |  |  |  |  |  |
| 1 | 40 | 11 | Normal | sEEG | (RAHC1-2)<br>1.5, 1.4 | (RPHG1-3)<br>0.9, 0.6, 0.4 | 1.2<br>[99.23%]<sup>a</sup> | 1 |
|  |  |  |  |  | (RAINS1-2)<br>2.7, 3.0 |  |  |  |
| 2 | 41 | 3 | Right hippocampal sclerosis | NA<sup>b</sup> | NA<sup>b</sup> |  | 4.3<br>[51.76%]<sup>a</sup> | 1 |
| 3 | 23 | 8 | Left hippocampal sclerosis | NA<sup>b</sup> | NA<sup>b</sup> |  | NA<sup>c</sup> | 1 |
| Neocortical temporal lobe epilepsy |  |  |  |  |  |  |  |  |
| 4 | 51 | 18 | Normal | sEEG | (LAMEG2-4)<br>1.6, 2.2, 2.3 | (LMEG4-7)<br>0.7, 0.4, 0.3, 0.6 | 0.8<br>[99.65%]<sup>a</sup> | NA<sup>d</sup> |
| 5 | 24 | 10 | Right temporal subcortical nodular heterotopia and schizencephalic cleft | sEEG | (RPHC7-9)<br>1.3, 0.6, 0.3 | (RPHG9)<br>1.4 | NA<sup>e</sup> | NA<sup>f</sup> |
| 6 | 19 | 18 | Right porencephalic cyst | ECoG | (GRID 28, 36)<br>0.3, 1.0<sup>g</sup> |  | 0.6<br>[99.95%]<sup>a</sup> | 1 |

NA = not applicable.
<sup>a</sup>Percentile of the beamformer power contrast at the location of the interictal dipole source solution. For example, 99% means that, at the location of the dipole, the beamformer power contrast is in the top 1% of its distribution.
<sup>b</sup>No iEEG recording.
<sup>c</sup>No interictal spikes.
<sup>d</sup>No surgical resection due to co-localization with eloquent language cortex.
<sup>e</sup>No valid interictal ECD source solution.
<sup>f</sup>No surgical resection (possible surgical approach under consideration).
<sup>g</sup>Estimate based on intraoperative photography.

### Intracranial EEG

Stereotactic EEG (sEEG) and electrocorticography (ECoG) were acquired from multi-contact depth or subdural electrodes, respectively, using standard clinical equipment and techniques. For sEEG, each electrode contact was mapped manually from the clinical computerized tomography/MRI reconstruction to the MRI scan in which the beamformer solution was originally obtained. For ECoG, electrode contact locations were reconstructed from intraoperative photographs.

### Ground truth determination

The definitive gold standard for identification of the SOZ in non-invasive source localization studies is iEEG,^2–4^ which, as described above, was not acquired in two patients, both of whom had unilateral MTLE based on clinical (video-EEG, neuropsychological assessment) and MRI investigations, and a postsurgical seizure-free outcome after anteromesial temporal resection (AMTR). A seizure-free outcome after epilepsy surgery is taken as evidence for correct identification of the “epileptogenic zone,” which conceptually includes the SOZ, but is not limited to the SOZ, and is thus a less exacting standard for the determination of spatial accuracy in source localization studies. Nevertheless, for this study, we included the two patients without iEEG because their specific, syndromic form of epilepsy (unilateral MTLE with ipsilateral hippocampal sclerosis) is so typical, and so uniformly associated with ipsilateral temporolimbic seizure onsets, that the combination of their clinical, MRI and surgical outcome data could be fairly considered to approximate a ground truth.

## Results

### Patient 1

One of the patient’s typical seizures was recorded, marked by a sense of déjà-vu and panic, evident in EEG over the right anterior temporal region, lasting 46 seconds, not readily apparent to visual analysis in MEG (Supplementary Fig. 1). MEG beamformer analysis showed two main ictal foci, one in the anterior hippocampus/parahippocampal gyrus and the other in the insula. Given normal brain MRI and the clinical and MEG features suggestive of insular involvement, unilateral sEEG was acquired from nine depth electrodes (Supplementary Fig. 2), documenting ictal onsets in the anterior hippocampus and adjacent parahippocampal gyrus, with seizure propagation to the insula (Supplementary Fig. 3). Fig. 1A-E shows a segment of rhythmic ictal iEEG activity, location of the maximally involved sEEG contacts, the MEG topoplots, and the ictal beamformer solutions. The distance from the mesial temporal beamformer focus to the hippocampal and parahippocampal sEEG contacts was within 1.5cm and 1cm, respectively (Table 1). The distance from the insula focus to the insular sEEG contacts was farther, 2.5-3cm, although the insular depth electrode implantation limited sEEG contacts to record mainly in the region of the anterior insula, which may have been slightly forward of the maximal region of propagated ictal activity, judging from the sEEG pattern and the beamformer localization in the mid-posterior insula.

**Figure 1.**
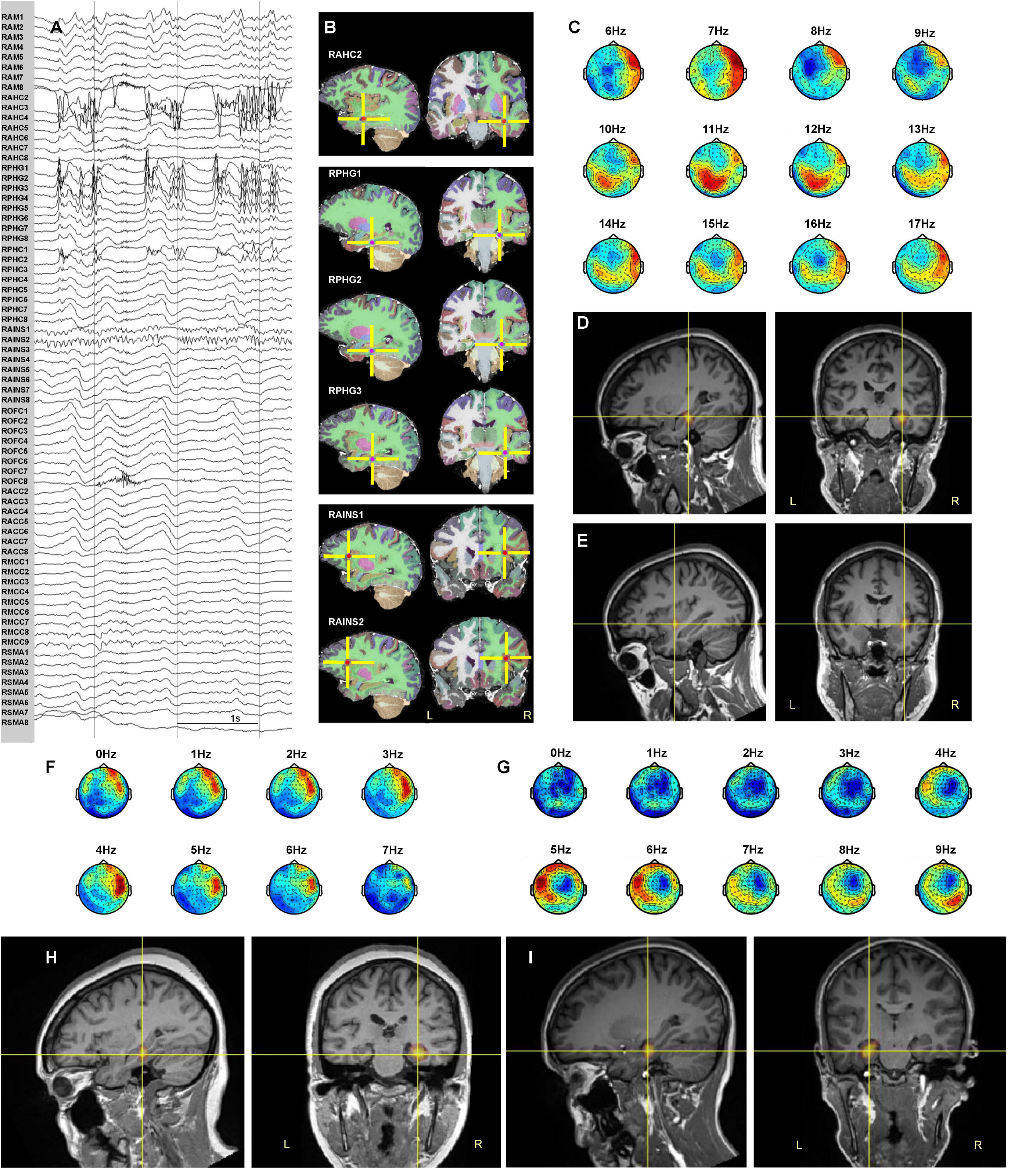
Beamformer localization of SOZ in MTLE patients. (A-E) Patient 1; (A) Ictal sEEG recording, late stage of seizure (see also Supplementary Fig. 3). Electrode labeling indicates target of deepest contacts (lowest numbers): AM = amygdala, AHC = anterior hippocampus, PHG = parahippocampal gyrus, PHC = posterior hippocampus, AINS = anterior insula, OFC = orbitofrontal cortex, ACC = anterior cingulate cortex, MCC = mid cingulate cortex, SMA = supplementary motor area. (B) Most active sEEG contact locations. (C) Topoplots corresponding to the most active ictal MEG frequencies. (D) Beamformer solution in the right parahippocampal gyrus. (E) Secondary beamformer solution in the right insula. (F) Topoplots corresponding to the most active ictal MEG frequencies in patient 2. (G) Topoplots corresponding to the most active ictal MEG frequencies in patient 3. (H) Ictal beamformer solution in the right hippocampus/parahippocampal gyrus, patient 2. (I) Ictal beamformer solution in the left parahippocampal gyrus, patient 3.

In addition, 139 right temporal interictal spikes were recorded, with ECD EMSI returning a right parahippocampal gyrus solution (Supplementary Fig. 4A) situated 1.2cm away from the mesial temporal ictal beamformer focus (Table 1). Comparing the ictal and interictal source solutions in a different fashion, one can assess how closely, at a percentile level, the interictal EMSI localization overlaps with the ictal BPC solution. In this case, the interictal EMSI solution was located within the top 1% of the ictal BPC distribution (Table 1).

A right AMTR resulted in a seizure-free outcome.

### Patient 2

Seven of the patient’s typical right temporal seizures were recorded, with no clinical manifestations apart from arousal from sleep in some events (Supplementary Fig. 5A). MEG beamformer analysis showed a right hippocampal/parahippocampal source (Fig. 1H).

The patient also had 64 right “classical” anterior temporal interictal spikes,^2,3^ localized by ECD EMSI to anterolateral temporal neocortex (Supplementary Fig. 4B), far (4.3cm) away from the mesial temporal ictal beamformer solution (Table 1).

Given MRI evidence of unilateral right hippocampal sclerosis and the ictal MEG localization, and with the knowledge that distant anterolateral temporal neocortical spikes are a common finding in MTLE,^2–4^ iEEG was not performed. A right AMTR resulted in seizure-freedom.

### Patient 3

Four subclinical electrographic seizures were recorded over the left temporal region, each lasting ~20 seconds (Supplementary Fig. 5B). The ictal MEG source reconstruction localized to the left parahippocampal gyrus using both DICS at 6Hz and unfiltered LCMV (Fig. 1I). DICS at 5Hz was associated with a secondary area of activation in the ipsilateral insula. No definite interictal epileptiform activity was recorded.

Given MRI evidence of unilateral left hippocampal sclerosis and the ictal MEG localization, iEEG was not performed. A left AMTR resulted in seizure-freedom.

### Patient 4

Brief left temporal subclinical seizures and rhythmic ictal-like bursts were recorded with an EEG frequency of approximately 10Hz. In addition, a single interictal spike focus was identified and modeled, with an EMSI ECD solution localized to the left superior temporal gyrus (Supplementary Fig. 4C). The MEG ictal topoplots suggested a very robust (same pattern across many frequencies) focal source, and the DICS beamformer solution at 10Hz showed a nearly identical location to the interictal EMSI solution (Table 1).

The sEEG implantation was partially guided by the MEG/EMSI solutions but also included additional bitemporal coverage (Supplementary Fig. 2). Ictal events were recorded maximally at sEEG contacts situated very close to the beamformer location (Fig. 2; Table 1).

**Figure 2.**
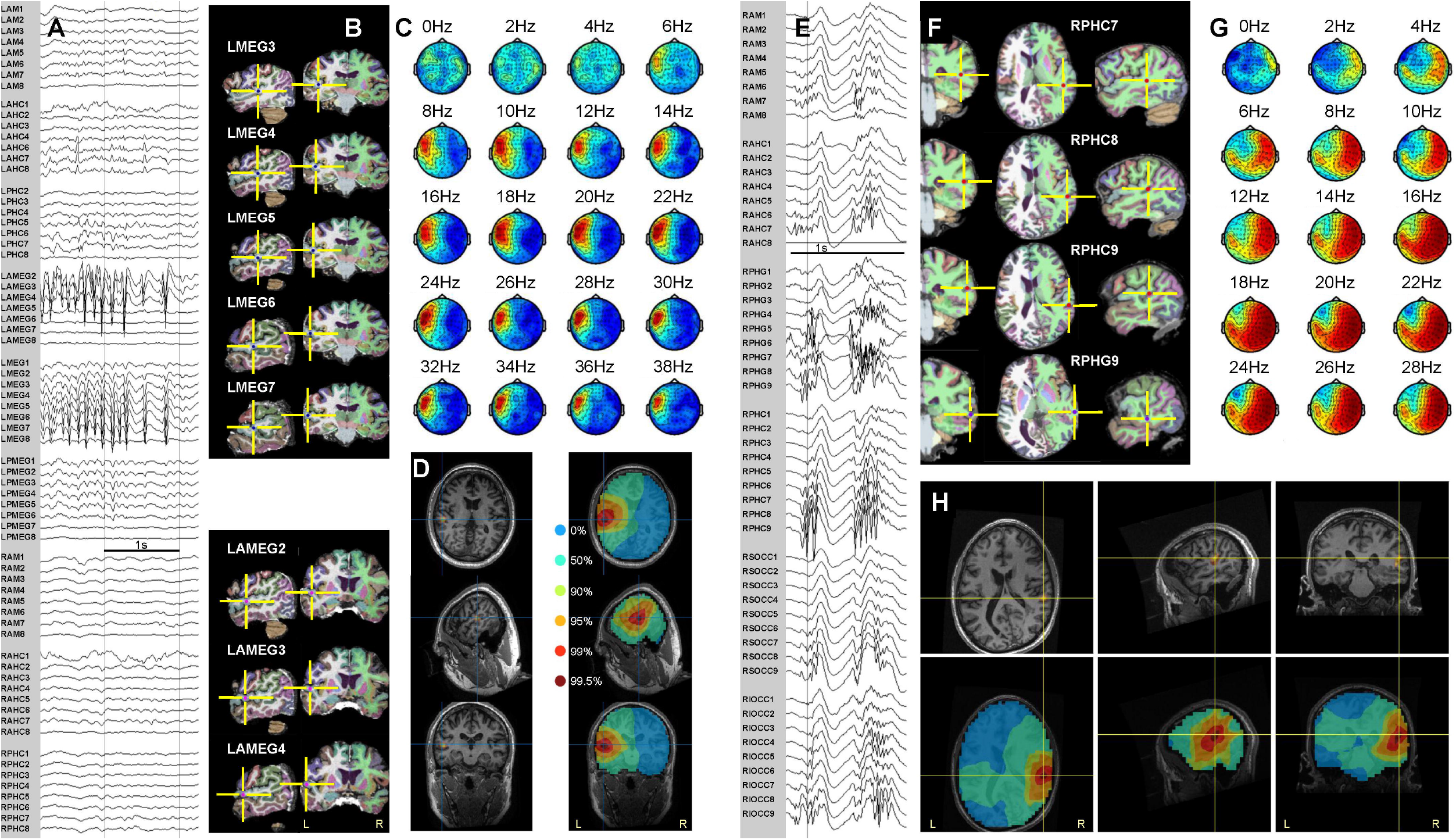
Ictal MEG beamformer localization and sEEG in temporal neocortical epilepsy patients. Patient 4 (A-D). Patient 5 (E-H). (A) Intracranial recording of a seizure. (B) Most active sEEG contact locations. (C) Topoplots corresponding to active ictal MEG frequencies. (D) Beamformer solution maximum in left superior temporal gyrus (left); unthresholded BPC by percentile levels (right). (E) Intracranial recording of a seizure. (F) Most active sEEG contact locations. (G) Topoplots corresponding to active ictal MEG frequencies. (H) Beamformer solution, thresholded (top) and unthresholded (bottom), in right middle temporal gyrus, superior to schizencephalic cleft. Intracranial electrode labeling, except for electrodes labeled MEG or OCC, indicates target of deepest contacts (lowest numbers): AM = amygdala, AHC = anterior hippocampus, PHG = parahippocampal gyrus, PHC = posterior hippocampus, AMEG = anterior MEG (targeting anterior to MEG beamformer solution), MEG = MEG beamformer solution (targeted directly), PMEG = posterior MEG (targeting posterior to MEG beamformer solution), OCC = occipital, S = superior, I = inferior, L = left, R = right.

### Patient 5

Multiple brief polyspike bursts were recorded, 0.2-3.5 seconds in duration, typically with a 20-30Hz beta maximum, occasionally evolving into a rhythmic ictal alpha frequency pattern maximal over the right posterior/basolateral temporal region. These rhythmic discharges were beamformed with DICS over the band 16-24Hz. No valid ECD solution could be obtained from modeling a complex, propagating interictal spike focus in the same region. Fifty-two sEEG contacts were implanted in the right hemisphere (Supplementary Fig. 2), partially guided by the ictal MEG solution. Particularly active sEEG contacts were located in the middle temporal gyrus, within 0.3-1.4cm of the beamformer solution (Fig. 2; Table 1).

### Patient 6

Abundant right posterior temporal rhythmic sharp wave activity was recorded. DICS and LCMV beamformer solutions using a variety of frequencies were quite robust, with occasional inferior mislocalizations ≤1cm into the cerebellum. We selected the DICS at 8Hz as a representative and plausible solution. Intraoperative ECoG confirmed that the beamformer solution was included in the area of the most active electrode contacts (Fig. 3), and surgical resection of the area resulted in seizure-freedom.

**Figure 3.**
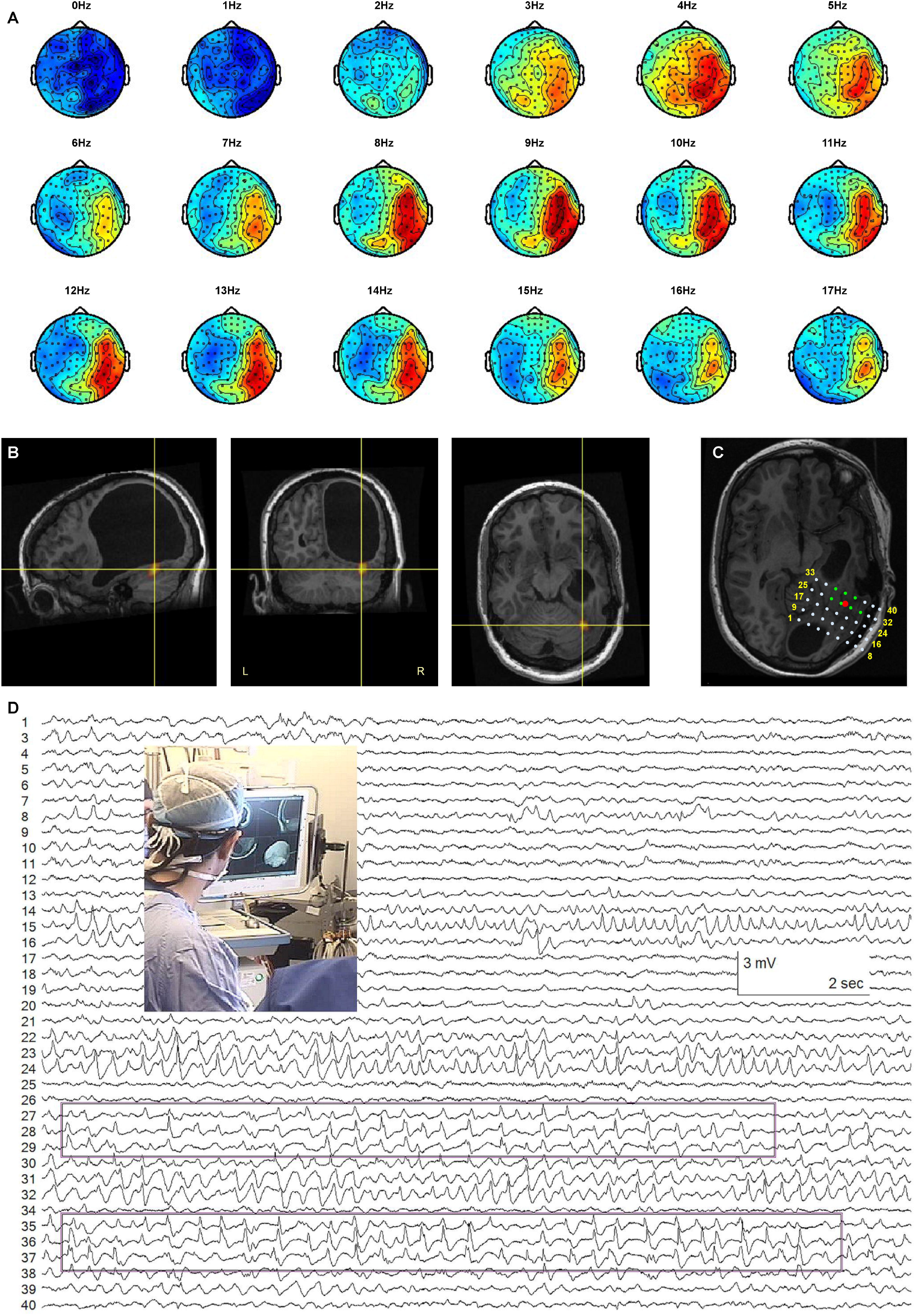
Ictal MEG beamformer localization and ECoG. Patient 6. (A) Topoplots corresponding to the rhythmic MEG sharp wave activity. (B) Beamformer source localization in basal temporal neocortex, below porencephalic cyst. (C) Representation of the position of the subdural ECoG grid. The contacts in green were most active during the intraoperative recording. The red dot is the position of the MEG beamformer solution. (D) Sample of ECoG recording with the most active contacts highlighted and a picture of the operating room setting where the beamformer solution is guiding the surgery.

EMSI of 227 right posterior temporal interictal spikes returned an ECD solution 0.6cm from the MEG ictal beamformer focus, at the 99.95 percentile level of the BPC solution (Table 1).

## Discussion

This report demonstrates the suitability of presurgical MEG beamformer analysis of ictal events in TLE, which can supersede or complement traditional analysis of interictal spikes. Beamforming also extends the number of events for analysis to include rhythmic sharp waves, polyspikes and small ictal-like bursts.^29,30^ Of particular relevance is the demonstration of robust seizure localization within deep structures in the MTLE patients. The ability to localize discretely hippocampal and parahippocampal gyrus seizures using MEG is, to our knowledge, something that has not been convincingly demonstrated previously, and is of obvious clinical importance.

The beamformer analysis method is proposed for use on any type of epileptiform activity, without any constraint on the activity’s temporal structure (unlike interictal spikes). As such, the ictal beamformer source reconstruction results may or may not localize to the same location as a patient’s interictal spike focus/foci. In MTLE, it is particularly common that the locations of interictal spike foci and a patient’s SOZ do not coincide.^2–4^ In the series of patients presented here we have shown instances where interictal and ictal analyses agreed in their localization, and one (MTLE) case where the interictal and ictal solutions were far apart. This is the nature of some epilepsies and the source localization results are in no way contradictory; instead, they simply represent the fact that the beamformer analysis is applied directly to ictal events. Another clear advantage of the method is in cases in which interictal spikes are absent in MEG-EEG recording, or too complex for traditional dipole or distributed source modeling (as in two of the patients in our series).

The ictal beamformer localizations showed good agreement with the gold standard iEEG. This is partly a consequence of surgical implantation planning having included the beamformer solutions, however, in all cases some of the most active iEEG contacts were located within 1cm of the BPC peak, a correlation that would not be achieved if the solutions were inaccurate. Moreover, iEEG was not restricted to the beamformer solution areas and typically covered a large brain volume.

It is necessary to acknowledge that a known limitation of beamforming is that solutions can be inaccurate if multiple sources are simultaneously active and correlated to some degree.^33^ Thus, in seizures suspected to be multifocal or widespread a note of caution should be added. In patient 1 two separate active foci were successfully discriminated, but this is not necessarily always possible.

In addition to the limitation in application to seizures that are focal in character (or multifocal with low correlation), there are assumptions regarding the spatio-temporal evolution of the ictal events that must be kept in mind. The method can miss the SOZ in cases with rapid ictal propagation to involve a larger area or multiple locations,^19^ something we have seen in a minority of extratemporal, frontal lobe cases similarly analyzed and contrasted with iEEG.

The natural trend in the development and application of diagnostic methods in epilepsy and EEG-MEG is toward an increased automatization and more independence from user input. However, in our proposal, and especially when intended for presurgical planning, it is key to integrate, rather than overlook, clinical expertise. Clinical guidance on a case-by-case basis is necessary from the initial phase of the analysis, when identifying and selecting events or identifying spatial and temporal differences in ictal features, to the last part of the analysis, evaluating the plausibility of the solution in the clinical context. In a majority of our ictal beamforming cases, temporal and extratemporal, MEG ictal source reconstruction results have seemed clinically plausible, correlating well with MRI abnormalities and/or focal semiological features. The detailed analyses of the TLE patients presented here, each with ground truth SOZ localizations, provide objective evidence to increase our confidence in ictal MEG source localization results acquired in the absence of ground truth. Nevertheless, although the beamformer tool we present is a valuable one, it is fallible, and its application is strengthened by clinical supervision.

## Acknowledgements

We are extremely grateful to Nat Shampur and his team of neurophysiology technologists for expert technical assistance. Victoria Barkley performed the intracranial electrode position reconstructions.

## Statement of Authorship

LGD and RW conceived the study and wrote the manuscript. LGD, AT and RW acquired and reviewed the EEG-MEG data and prepared the figures. LGD originated the ictal beamforming method and performed the beamforming analyses. Interictal EMSI analyses were done by RW. TV performed the iEEG electrode implantations and surgical resections. All authors critically reviewed the manuscript and approved the final version.

## Disclosures

The authors have no conflicts of interest to declare.

## Data availability

Individual deidentified MEG-EEG data files used for the analyses in this study will be made available upon reasonable request to the authors.

## Funding

None.

## Notes

### Competing Interest Statement

The authors have declared no competing interest.

