## Supplementary Fig. for "Beamforming seizures from the temporal lobe"

Beamforming seizures from the temporal lobe. Supplementary Figures 1-5.

**Supplementary Figure 1.**

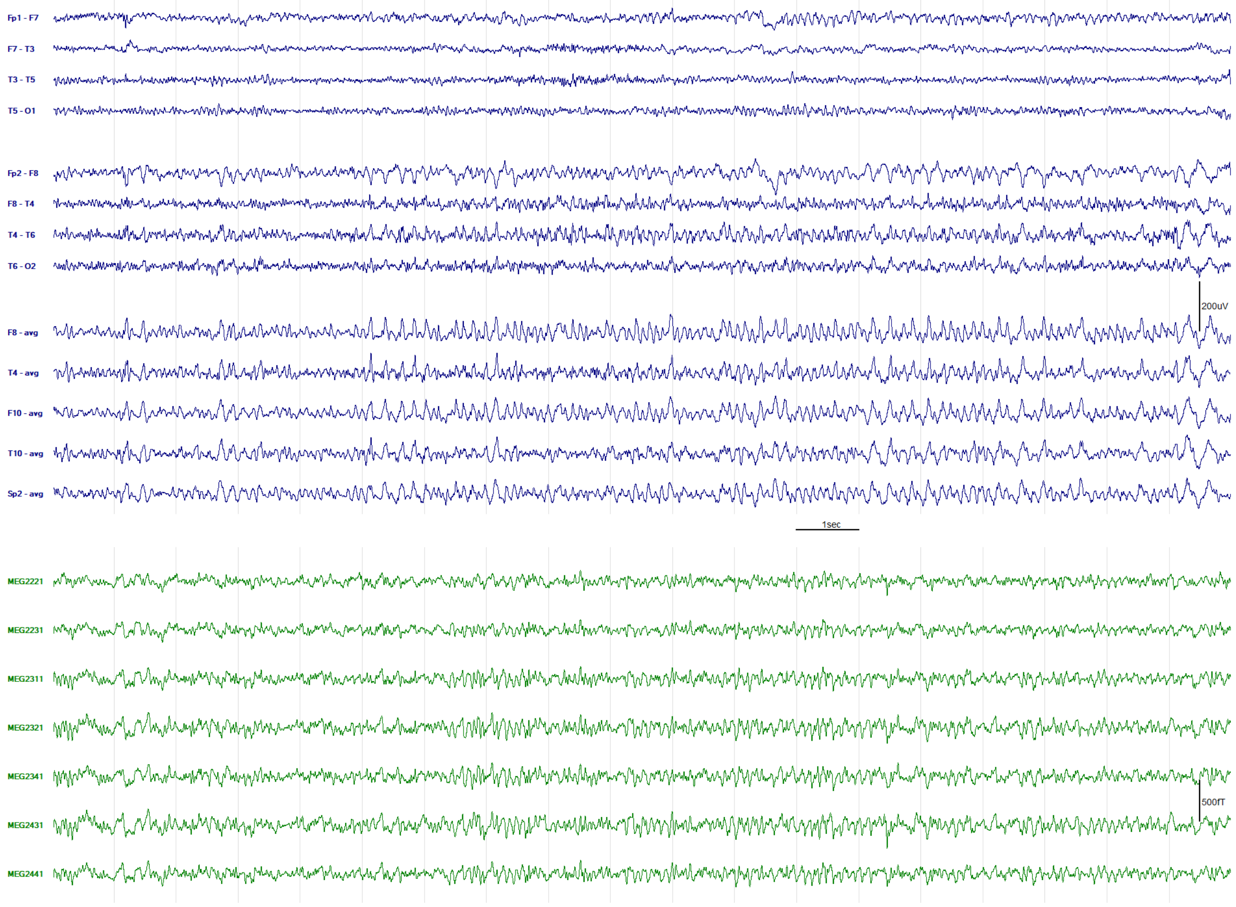


EEG-MEG recording of seizure onset in patient 1, the rhythmic ictal activity appearing maximal in EEG at F8, F10, Sp2 > T4, T10. The ictal discharge was not easily identifiable in MEG: seven magnetometer channels that appeared to show an ictal correlate to the seizure evident in EEG are shown. Filter bandpass for figure = 1-30Hz; ave = common average reference.

**Supplementary Figure 2.**

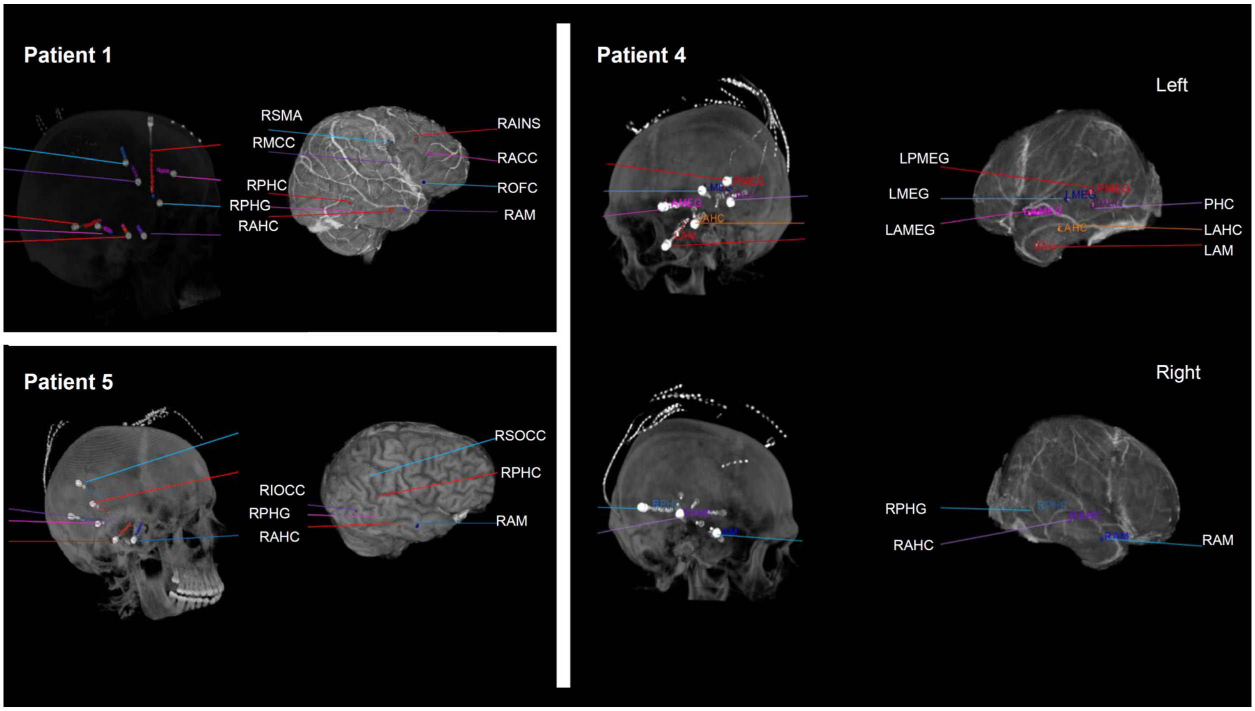


Intracranial sEEG depth electrode implantations; patients 1, 4 and 5. Electrode labels described in figure legends of main text.

**Supplementary Figure 3.**

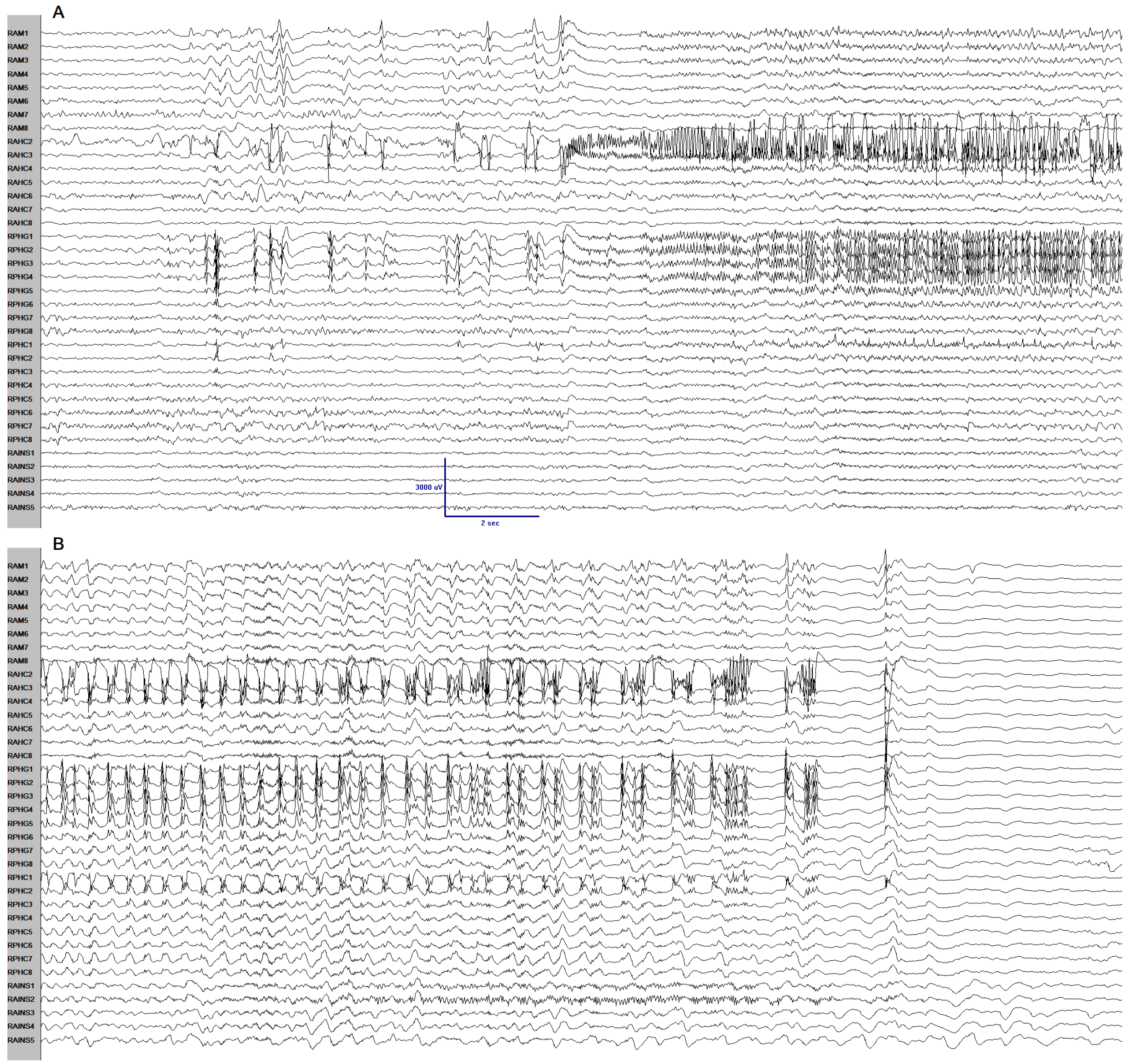


Ictal event recorded with sEEG in patient 1. (A) Seizure onset in anterior hippocampus (RAHC2) and parahippocampal gyrus (RPHG1-4). (B) Ictal propagation to posterior hippocampus (RPHC1) and, independently, anterior insula (RAINS1-2). Filter bandpass for figure = 1-50Hz.

**Supplementary Figure 4.**

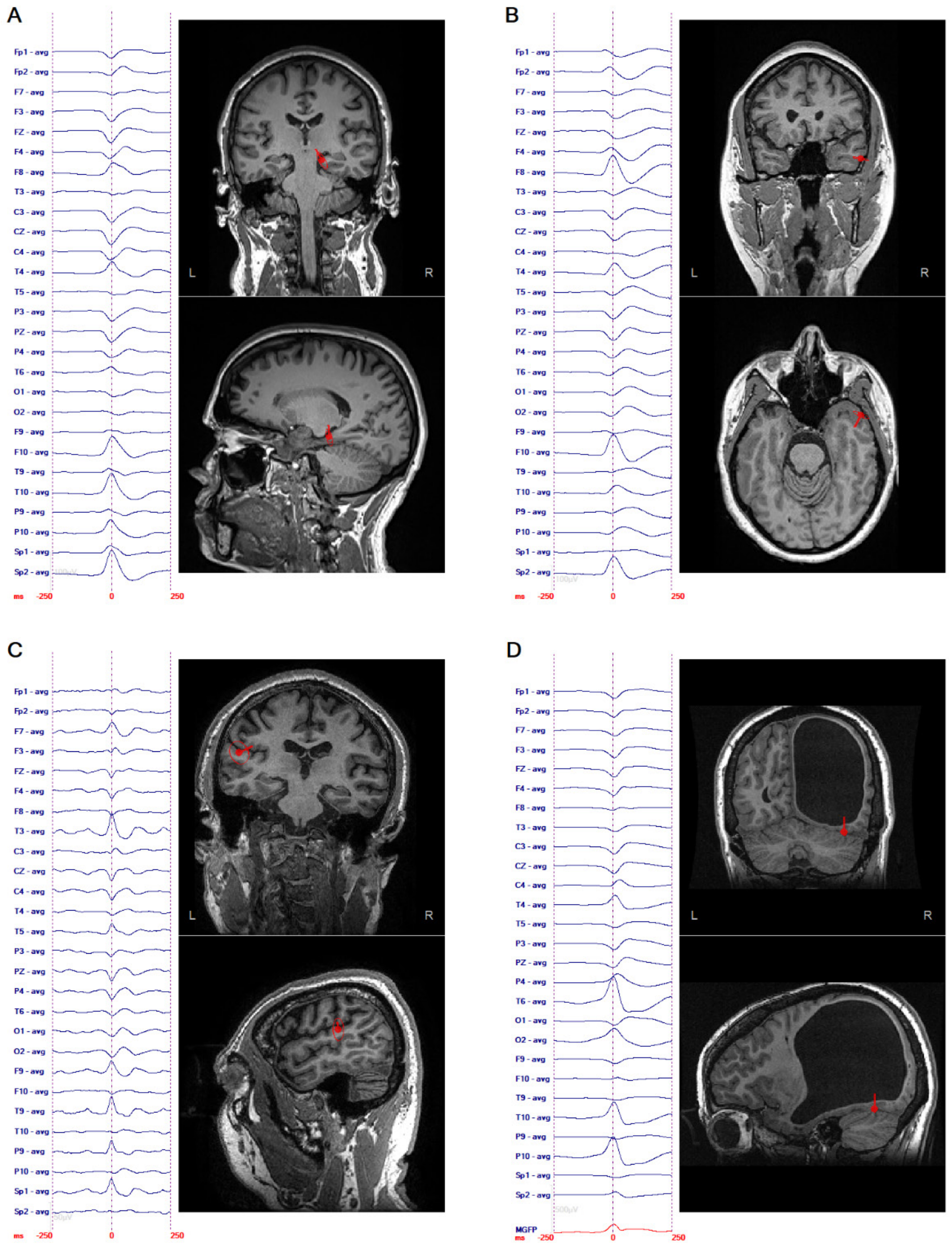


EMSI of interictal spike foci in patients 1 (A), 2 (B), 4 (C) and 6 (D). ECD modeling performed as described previously[1-3] on averaged spikes (patient 1, n=139; patient 2, n=64; patient 4, n=25; patient 6, n=227). Filter bandpass for figure = 1-70Hz; bandpass for ECD modeling = 1-30Hz.

**Supplementary Figure 5.**

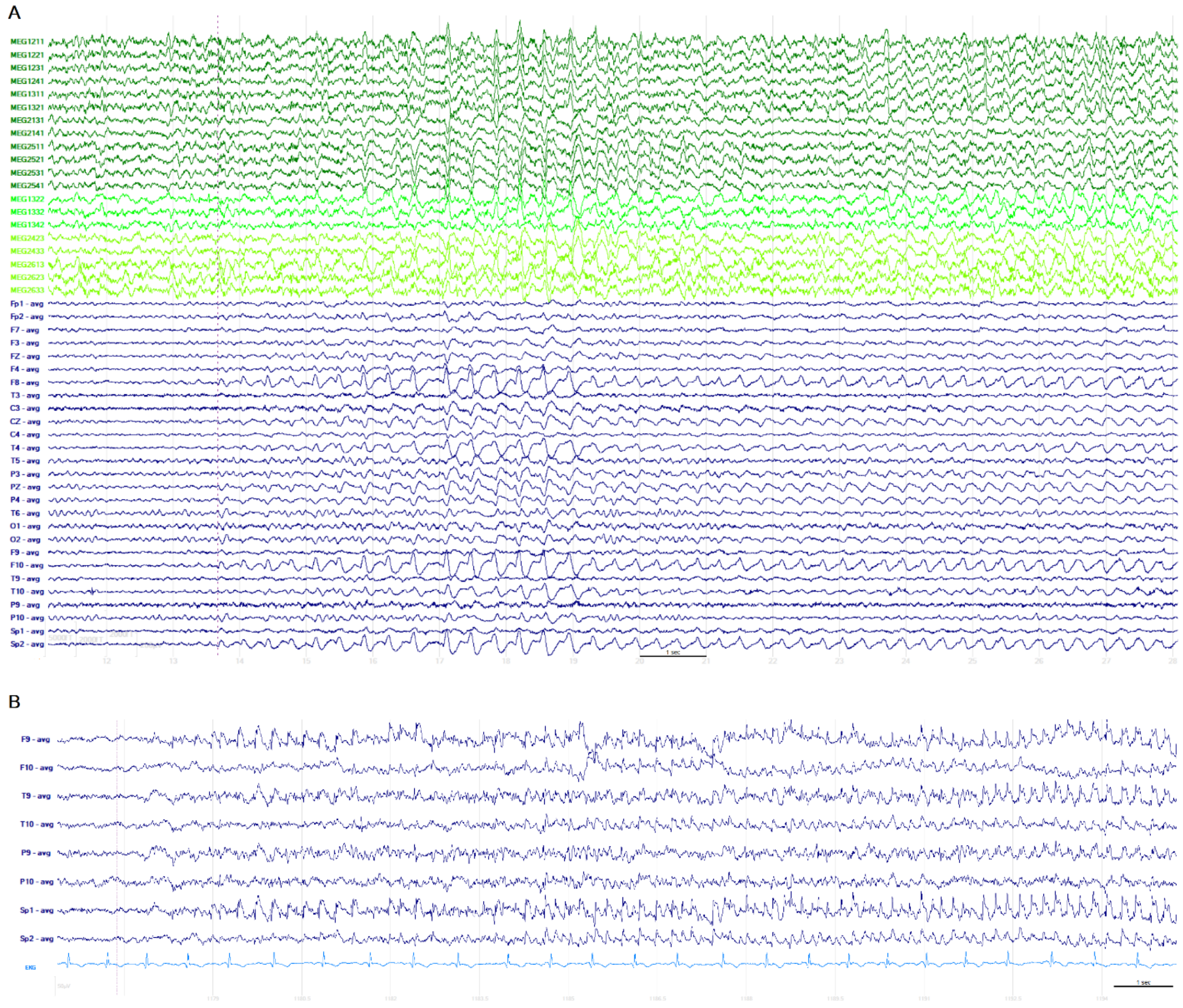


(A) EEG-MEG recording of seizure onset in patient 2, the rhythmic ictal activity appearing maximal in EEG at F8, F10, Sp2, simultaneously apparent in a selection of magnetometer (dark green) and orthogonal planar gradiometer (green and light green) MEG channels. (B) EEG ictal onset of seizure during EEG-MEG recording in patient 3, appearing maximal at F9, Sp1 > T9. The ictal discharge was not visually identifiable in MEG. Filter bandpass for figures = 1-70Hz; ave = common average reference.
